# Genome Report: A chromosome-level genome assembly of the varied leaved jewelflower, *Streptanthus diversifolius,* reveals a recent whole genome duplication

**DOI:** 10.1101/2024.05.20.595032

**Authors:** John T. Davis, Qionghou Li, Christopher J. Grassa, Matthew Davis, Sharon Y. Strauss, Jennifer R. Gremer, Loren H. Rieseberg, Julin N. Maloof

## Abstract

The Streptanthoid complex, a clade of primarily *Streptanthus* and *Caulanthus* genera in the Thelypodieae tribe (Brassicaceae) is an emerging model system for ecological and evolutionary studies. This Complex spans the full range of the California Floristic Province including desert, foothill, and mountain environments. The ability of these related species to radiate into dramatically different environments makes them a desirable study subject for exploring how plant species expand their ranges and adapt to new environments over time. Ecological and evolutionary studies for this complex have revealed fascinating variation in serpentine soil adaptation, defense compounds, germination, flowering, and life history strategies. Until now a lack of available genomic resources has hindered the ability to relate these phenotypic observations to their underlying genetic and molecular mechanisms. To help remedy this situation we present here a chromosome-level genome assembly of *Streptanthus diversifolius*, a member of the Streptanthoid Complex, developed using Illumina, Hi-C, and HiFi sequencing technologies. Construction of this assembly also provides further evidence to support the previously reported recent whole genome duplication unique to the Thelypodieae tribe. This whole genome duplication may have provided individuals in the Streptanthoid Complex the genetic arsenal to rapidly radiate throughout the California Floristic Province and to occupy commonly inhospitable environments including serpentine soils.

## Introduction

The California Floristic Province (CFP) is a region comprising much of the state of California west of the Sierra Nevada Mountains, and includes portions of Oregon and Baja California (Howell, 1957). It is a hotspot for biodiversity due to its Mediterranean-type climate with warm dry summers and cool wet winters. The CFP is home to more than 700 genera and more than 4000 vascular plants native to California. Among this collection is the ‘Streptanthoid Complex’ which is a set of ca. 60 taxa belonging to the Thelypodieae tribe (Burrell, 2010; Burrell et al., 2011; Burrell and Pepper, 2006). This complex is comprised primarily of the *Streptanthus* and *Caulanthus* genera, and while the members are genetically closely related, they occupy a large range of environments including the deserts, mountains, and foothills of the CFP (Al-Shehbaz, 2010).

*Streptanthus* is well known for extreme edaphic specialization, particularly to infertile serpentine soils. Serpentine is a unique soil known for its extraordinarily low Ca to Mg ratio, low levels of micronutrients, elevated heavy metal concentrations, and poor water retention (Brady et al., 2005; Kruckeberg, 2006). Plants grow best when the Ca to Mg ratio of the soil is close to one or greater (Palm and Van Volkenburgh, 2014) which makes the low Ca to Mg ratio of particular interest as Mg^2+^ concentrations in serpentine soils can exceed those of Ca^2+^ by almost tenfold (Alexander, 2007). This imbalance of alkaline-earth metals has a profound effect on plant ecology and evolution, often supporting unique plant communities of serpentine tolerant species, as well as serpentine endemics that occur on no other soil type. Serpentine soil usage has four or five independent origins in the Streptanthoid Complex (Cacho and Strauss, 2014) and has led to more than half of the ∼35 known species in the complex to be endemic to serpentine soils (Al-Shehbaz, 2010; Baldwin et al., 2012; Safford et al., 2005). The genus has been the subject of several classic studies on the evolution of edaphic specialization and is an emerging model system for understanding the evolution of edaphic specialization in herbaceous plants. *Streptanthus* has also been the subject of molecular phylogenetic investigations, providing a robust phylogenetic context in which to examine the evolution of specific traits during diversification(Cacho et al., 2021; Cacho and Strauss, 2014; Weber et al., 2018).

To improve genomic resources for research in *Streptanthus* and its allies, we carried out whole-genome shotgun sequencing, Hi-C sequencing, and PacBio HiFi sequencing on the nuclear genome of the varied leaved jewelflower, *S. diversifolius.* While *S. diversifolius* is not found on serpentine soils, it has the smallest genome known in the genus (0.36 GB), and is a known diploid (2n=28) (Cacho et al., 2021). A reference genome for *S. diversifolius* will allow for gene discovery, genetic mapping, and re-sequencing of other *Streptanthus* species, all in support of ongoing studies. The reference genome sequence described here will allow mechanistic knowledge from the related model genera Arabidopsis and Brassica to be leveraged for understanding the molecular basis of serpentine specialization, heavy metal tolerance and hyperaccumulation, and climate adaptation. Such insights may lead to improvements of closely related crops in the Brassicaceae family, as well as industrial applications in phytoremediation and phytomining. Additionally, a greater understanding of the *Streptanthus* clade will allow us to build upon previously work including the evolution of glucosinolate defense (Cacho et al., 2015) and the seasonal germination niche (Gremer et al., 2020; Worthy et al. submitted). Combined these analyses will assist in the conservation and management of these species, many of whom are currently endangered or threatened.

## Methods & Materials

### Plant collection, DNA isolation, and Sequencing

Plant collection, DNA isolation, and sequencing occurred in three different rounds. In the first round, several whole, flowering plants of *S. diversifolius* were collected in April 2013 from a naturally occurring population on Table Mountain, Butte County, California (D. O. Burge 1389; voucher deposited at DAV). Tissues of these plants were dried on silica gel desiccant at room temperature before being returned to the lab at the University of British Columbia. Total genomic DNA was extracted from the leaves, stems, and unopened flower buds of a single plant using the Qiagen (Limburg, Netherlands) DNeasy Kit according to the manufacturer’s instructions. A total of 24 extractions were performed. Pooled extractions were then concentrated by adding 1/10 volume of 3M sodium acetate, pH 5.2, and 2 volumes of of −20°C 95% EtOH. After centrifugation, the DNA pellet was dried and resuspended in pure, nuclease-free water. Aliquots of the same DNA preparation were then used to construct four sequencing libraries according to the manufacturer’s protocols (Illumina, http://www.illumina.com) at the Innovation Centre, Genome Quebec. Library insert sizes were chosen to be compatible with the Allpaths-LG genome assembler (Gnerre et al., 2011). We prepared two libraries designed to overlap when sequenced as paired ends: a 180 bp library sequenced on an Illumina HiSeq with 2×100bp reads, and a 450 bp library sequenced on an Illumina MiSeq with 2×300bp reads. We also prepared two mate-pair libraries with insert sized of approximately 4700 bp and 8200 bp.

In the second round, seeds from *S. diversifolius* were collected in 2016 from a naturally occurring population on Table Mountain, Butte County, California. It should be noted that this is a separate seed collection from that described above. Eight cones containing a mixture of 50% Ron’s Mix and 50% sand were saturated with nutrient water. A small divot was then made in each cone where 3-4 seeds were placed before being covered with a small amount of the soil mixture. The cones were then placed in a rack and covered with plastic wrap to prevent the top layer of the soil from drying out. The covered rack was then placed in a growth chamber set to 22*C with a light/dark cycle of 12/12. Two weeks after being placed in the growth chamber, the plastic wrap was removed and the recently germinated seedlings were exposed to the air. The seedlings remained in the growth chamber with nutrient water being provided every other day. Once the seedlings on average had attained approximately 8-10 true leaves, two plants were randomly selected for Hi-C sequencing. Young leaves from each plant were collected separately until approximately 0.5 grams were obtained. The leaves from each plant were than processed separately using Phase Genomics’ Proximo Hi-C Kit version 3.0 following the standard plant protocol. The libraries where then sent to the University of California, Davis Genome Center where 150bp paired-end sequencing was performed using an Illumina NovaSeq 6000. A total of ∼240 million read pairs were sequenced resulting in ∼200X coverage of the genome.

In the third round, leaf tissue from 3 different *S. diversifolius* individuals located at Table Mountain, Butte Country, CA were harvested individually in May 2023. The 3 samples weighing 0.66g, 0.59g, and 0.52g were placed on ice and transported to UC Davis where they were flash frozen in liquid nitrogen upon arrival. All 3 samples were then sent to the UC Davis genome center for HMW DNA extraction. DNA was successfully extracted from the 0.66g sample and used for sequencing and the remaining 2 leaf samples were kept for backup. Sequencing was also performed at the UC Davis genome center on PacBio’s Revio sequencing platform. The reads were then processed using SMRT Link version 12.0.0.177059 resulting in 4,689,053 HiFi reads with a mean read length of 11,052 base pairs and mean read quality of 30. The HiFi yield of 51.8 Gb represents a genome coverage of ∼144X.

### Genome Assembly

Starting with the Illumina short reads from round one, raw sequence reads were filtered and trimmed of their adapter sequences using Trimmomatic (Bolger et al., 2014). Trimmed and filtered reads were assembled using Allpaths-LG with the haploidify=T parameter. Biological contaminants in the resulting assembly were identified against the NCBI NT database using blastn megablast (Sayers et al., 2022). Seven scaffolds with a match of at least 85% and at least 300 bp to a database sequence belonging to non-plant taxa were removed from the Allpaths-LG assembly. Artificial contaminants were identified using NCBI’s VecScreen protocol. Contaminants at scaffold ends were removed; artifacts within scaffolds were masked.

To improve the contiguity of the assembly from round one, the Hi-C sequencing data from round two was added to the assembly. The Hi-C reads were trimmed using Trimmomatic version 0.39 in paired end mode with the parameters ILLUMINACLIP:adapters.fa:2:30:10 LEADING:3 TRAILING:3 SLIDINGWINDOW:4:15 MINLEN:36 resulting in ∼180 million surviving read pairs. The reads were then mapped to round one scaffolds using the Arima-HiC Mapping Pipeline (Arima Genomics, n.d.). Following mapping, the alignment file was sorted by read name using Samtools (Li et al., 2009) sort. The sorted alignment file along with the scaffolds were input into the Hi-C scaffolding program YaHS (Zhou et al., 2022).

The Hi-C scaffolded assembly showed signs of a possible whole genome duplication along with highly probable mis-joins of the contigs. To investigate these two occurrences, a new genome assembly was created on the Jetstream2 cloud computational platform (Hancock et al., 2021) using the HiFi sequencing data from round three. The HiFi reads were converted from BAM format to FASTQ format for assembly. The FASTQ reads were assembled using the program HiFiasm version 0.19.5-r593 (Cheng et al., 2021) using default parameters with the –primary flag selected. The genome assembly created using HiFiasm was used for future analyses presented here.

### Repeat and gene annotation

The assembly was annotated using the MAKER pipeline (Cantarel et al., 2008; Holt and Yandell, 2011). First a custom repeat library for *S. diversifolius* was made using the Maker-P pipeline (Campbell et al., 2014b, 2014a). Augustus (Keller et al., 2011) retraining parameters were also calculated using BUSCO v4.1.4 (Manni et al., 2021) in long and genome mode and the brassicales_odb10 lineage dataset. The assembly was input into Maker v3.01.04 along with the repeat library, Augustus retraining parameters, and transcripts from six *Streptanthus* clade species. A second round of Maker was performed using same inputs except this time a newly made *S. diversifolius* hmm file made using Snap (Korf, 2004) v2006-07-28 and the GFF file created in the first round of Maker were included. In the second round of Maker the parameter of AED=0.5 was also set, previously AED=1, to remove transcripts which had weak evidence support. A total of 40,606 gene models were created. These gene models were post processed following the MAKER Tutorial for WGS Assembly and Annotation Winter School 2018 (https://weatherby.genetics.utah.edu/MAKER/wiki/index.php/MAKER_Tutorial_for_WGS_Assembly_and_Annotation_Winter_School_2018) where they were aligned to the UniProt database using blastp and processed using interproscan (Jones et al., 2014). These results were then incorporated into the final annotations.

### Telomere annotation

To test whether we had captured telomeres, we used the Biostrings (Pagès et al., 2024) package to count occurrences of the 21bp sequence “TTTAGGGTTTAGGGTTTAGGG” and its reverse complement (representing 3 repeats of the canonical plant telomere repeat sequence) in 1,000,000 base windows across the genome. We retained windows that had at least 20 copies of the query sequence.

### Alignment to *A. thaliana*

To assess the contiguity of the Hi-C and HiFi draft assemblies, the *S. diversifolius* genomes were aligned to a reference *A. thaliana* genome (TAIR 10)(Berardini et al., 2015). The genomes were aligned in protein space using the promer alignment tool contained within the mummer software package (Marçais et al., 2018). Promer was run using the ‘maxmatch’ flag and the default setting for the other parameters. Following alignment, the delta files were then filtered using delta-filter using minimum alignment length and sequence identity cutoffs of 1000 bp and 85% respectively. The filtered delta files were then plotted using mummerplot. Points in the plots were colored based on the ancestral crucifer karyotype (ACK) blocks (Lysak et al., 2016) defined within *A. thaliana*.

### Genomic data collection

To investigate the possibility of a whole genome duplication, comparative genomic analyses were performed. The genome data and annotation files of following Brassicaceae species were obtained from public repositories: *Caulanthus amplexicaulis*, *Stanleya pinnata*, *Arabidopsis thaliana* (Cheng et al., 2017), *Arabidopsis lyrata* (Hu et al., 2011) from Phytozome (https://phytozome-next.jgi.doe.gov/) (Goodstein et al., 2012); *Cardamine hirsute* (Gan et al., 2016) from *Cardamine hirsuta* (http://chi.mpipz.mpg.de/index.html); *Euclidium syriacum* was from NCBI accession number (GCA_900116095.1); *Draba nivalis* (Nowak et al., 2021) from Dryad (https://datadryad.org); *Arabis alpina* (Jiao et al., 2017)w from (http://www.arabis-alpina.org/index.html); *Thlaspi arvense* genome (Geng et al., 2021) from (http://pennycress.umn.edu/); *Schrenkiella parvula* e (Dassanayake et al., 2011) from thellungiella.org; *Brassica oleracea* from Genoscope (http://www.genoscope.cns.fr/externe/plants/chromosomes.html); *Raphanus raphanistrum* ssp. *raphanistrum* (Zhang et al., 2021) from (https://ngdc.cncb.ac.cn/gwh/#) under accession number PRJCA003033; and *Aethionema arabicum* e (Fernandez-Pozo et al., 2021) from (https://plantcode.cup.uni-freiburg.de/aetar_db/downloads.php)

### Species tree construction

To construct a species tree, we identified single-copy orthogroups from the proteins of the 14 genomes listed above, including *S. diversifolius*, using OrthoFinder (Emms and Kelly, 2019). This identified 118 single-copy orthogroups that were then aligned using MAFFT (Katoh and Standley, 2013). Pal2nal was used to perform codon alignment (Suyama et al., 2006). TrimAl was used to trim poorly aligned segments from the alignments (Capella-Gutiérrez et al., 2009). Based on high-quality alignments, the phylogenetic trees for the 118 single-copy orthogroups were constructed using the IQ-TREE (Minh et al., 2020). Finally, the species tree was inferred using ASTRAL-III (Zhang et al., 2018).

### Identification of whole genome duplication

To identify whole-genome duplication events in *Streptanthus*, we compared the *S. diversifolius* genome with 13 other genomes. Initially, the DupGene_Finder pipeline (Qiao et al., 2019) was used to identify WGD genes across the 14 genomes, including *S. diversifolius*, while removing tandem genes. Subsequently, the Ka_Ks_pipeline (Qiao et al., 2019) was utilized to calculate the number of substitutions per synonymous site (Ks) values for WGD pairs. We made some modifications to the previous Ks fitting process (identify_Ks_peaks_by_fitting_GMM) to allow fitting based on the Ks values of gene pairs. To place the WGD events, based on the phylogenetic relationships of the 14 species, we identified orthologs between *S. diversifolius* and *C. amplexicaulis* and orthologs between *S. diversifolius* and *S. pinnata*, calculated the Ks values for these orthologs, and plotted the Ks distribution. Additionally, we used MCscan (Tang et al., 2008) (python version, https://github.com/tanghaibao/jcvi) to create dot-plots for *S. diversifolius* vs *S. diversifolius* and *S. diversifolius* vs *A. thaliana* to further confirm the recent WGD events. The timing of the WGD events was estimated to be between 11.54 to 14.06 million years ago (MYA), based on the formula T=Ks/2r (r=7.1 × 10−9 ± 0.7 × 10−9) (Ossowski et al., 2010).

### Sub-genome differentiation

Two methods were used to look for possible differentiation among sub-genomes. First, SubPhaser (Jia et al., 2022) was run with default parameters (k = 15, q = 200) to look for evidence of unique kmers among homeologs. Second, the CoGe framework (Lyons and Freeling, 2008) was used to align *S. diversifolius* coding regions against those from *Arabidopsis thaliana* and to perform a fractionation bias (Joyce et al., 2017) analysis. A syntenic ratio of 2:1 *S. Diversifolious* : *A. thaliana* was specified for Quota Align (Tang et al., 2011); otherwise, default parameters were used.

### Gene family evolution

To identify expanded and contracted gene families, the CAFE5 (Mendes et al., 2021) software was utilized. Following the CAFE5 documentation, orthogroups previously identified by OrthoFinder with more than 100 gene copies in one or more species were excluded in downstream analysis. The previously described species tree was transformed into an ultrametric tree, using MCMCtree which is incorporated in PAML v4.10.7 (Yang, 2007), utilizing the age of the most recent common ancestor of *A. arabaicum* and the remaining 13 other species as 35.2 million years ago (MYA), based on a previous study (Nowak et al., 2021). The ultrametric tree and the filtered orthogroups were then used as input for CAFE5. Testing was perfomed on different -k parameters to determine the optimal K value based on the Model Gamma Final Likelihood value, where the best -k value was found to be 2. Finally, significantly expanded and contracted families were extracted from the result files.

### Positive selection tests

The identification of positively selected genes (PSGs) in Streptanthus was conducted using the branch-site model in codeML implemented in PAML v4.10.7 (Yang, 2007). Due to a relatively small number of single-copy orthogroups (only 118 single-copy orthogroups), low-copy genes were used to identify PSGs. Initially, orthogroups with less than three copies across all species were identified based on OrthoFinder results. Codon alignments were then performed using the OMM_MACSE pipeline within MACSE v11.05 (Ranwez et al., 2018), which employs MAFFT (Katoh and Standley, 2013) and PRANK (Löytynoja, 2014) for codon alignment. Based on codon alignments from the OMM_MACSE pipeline, GWideCodeML v1.1 (Macías et al., 2020) was used for genome-wide PSG identifications. Briefly, using the species tree, Streptanthus was set as the foreground branch. Sites showing significant positive selection were identified by comparing the likelihood ratio test (LRT) values between the alternative model and the null model. GWideCodeML automates these steps and the comparison, ultimately identifying significantly selected sites and genes. A gene is considered a PSG only if it has an p-value less than 0.05 and also has significantly selected sites.

## Results

The construction of this new *S. diversifolius* whole genome assembly took place over multiple rounds, with each round producing an improved assembly compared to its predecessor. Starting with an Illumina short read assembly and progressing to a HiFi assembly, the results of each iteration are described below. Following the assembly are the results of analyses investigating the presence of a suspected whole genome duplication and positive selection of gene families.

### Round 1: Illumina Assembly

The four Illumina sequencing libraries, HiSeq with 2×100bp reads, MiSeq with 2×300bp reads, and mate-pair libraries with insert sizes of approximately 4700 bp and 8200 bp were used for the initial assembly. Assembly was performed using the whole-genome shotgun assembler ALLPATHS-LG and subsequently postprocessed using NCBI’s VecScreen protocol. The resulting assembly had a total length of 314 Mb, a scaffold N50 of 470 Kb, and was comprised of 4,627 scaffolds (Table 1). Genome completeness of the assembly using BUSCO version 5.5.0 and the embryophyta odb10 database found the assembly to have a complete BUSCO percentage of 98.7% with an approximate even split between single-copy and duplicate BUSCOs (Table 2).

**Table 1.**
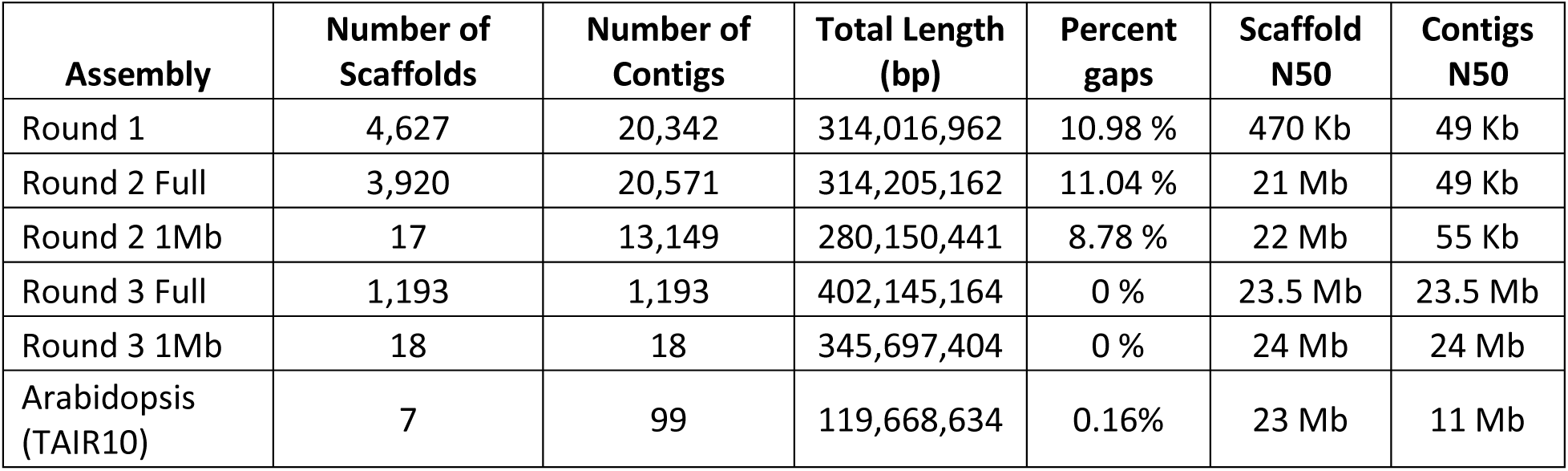
Assembly statistics for each round of assembly.

| Assembly | Number of Scaffolds | Number of Contigs | Total Length (bp) | Percent gaps | Scaffold N50 | Contigs N50 |
| --- | --- | --- | --- | --- | --- | --- |
| Round 1 | 4,627 | 20,342 | 314,016,962 | 10.98 % | 470 Kb | 49 Kb |
| Round 2 Full | 3,920 | 20,571 | 314,205,162 | 11.04 % | 21 Mb | 49 Kb |
| Round 2 1Mb | 17 | 13,149 | 280,150,441 | 8.78 % | 22 Mb | 55 Kb |
| Round 3 Full | 1,193 | 1,193 | 402,145,164 | 0 % | 23.5 Mb | 23.5 Mb |
| Round 3 1Mb | 18 | 18 | 345,697,404 | 0 % | 24 Mb | 24 Mb |
| Arabidopsis (TAIR10) | 7 | 99 | 119,668,634 | 0.16% | 23 Mb | 11 Mb |

**Table 2.**
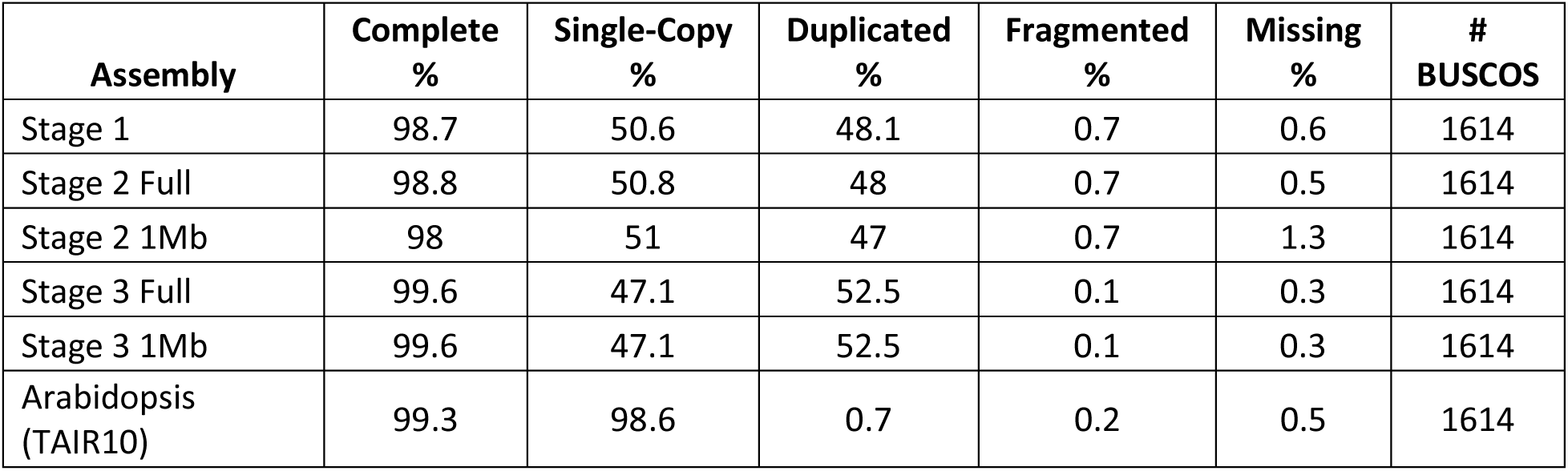
BUSCO summary statistics. Analysis completed using BUSCO version 5.5.0 in genome mode and the embryophyta odb10 dataset.

| Assembly | Complete % | Single-Copy % | Duplicated % | Fragmented % | Missing % | # BUSCOS |
| --- | --- | --- | --- | --- | --- | --- |
| Stage 1 | 98.7 | 50.6 | 48.1 | 0.7 | 0.6 | 1614 |
| Stage 2 Full | 98.8 | 50.8 | 48 | 0.7 | 0.5 | 1614 |
| Stage 2 1Mb | 98 | 51 | 47 | 0.7 | 1.3 | 1614 |
| Stage 3 Full | 99.6 | 47.1 | 52.5 | 0.1 | 0.3 | 1614 |
| Stage 3 1Mb | 99.6 | 47.1 | 52.5 | 0.1 | 0.3 | 1614 |
| Arabidopsis (TAIR10) | 99.3 | 98.6 | 0.7 | 0.2 | 0.5 | 1614 |

### Round 2: Hi-C Assembly

Looking to improve the contiguity of the Illumina assembly, two Hi-C sequencing libraries were prepared using tissue from two separate *S. diversifolius* plants whose seeds came from a population in the same geographic location as the original plant samples. Both sequencing libraries were aligned to Illumina assembly using the Aria HiC Mapping Pipeline and the resulting alignment files were input into the Hi-C scaffolding program YaHS. The scaffolded assembly showed signs of improvement compared to the original Illumina assembly. The scaffolding resulted in the overall size of the assembly increasing slightly due to the introduction of scaffolding gaps. The main improvements were a decrease in the number of scaffolds from 4,627 to 3,920 scaffolds and an increase in the scaffold N50 from 470 Kb to 21Mb. Additionally, 17 scaffolds were greater than 1Mb in length and encompassed 89% of the total assembly length (Table 1). Scaffolding did not have a significant effect on BUSCO composition and the majority of the BUSCOs were found to be contained in the 17 previously mentioned scaffolds (Table 2).

To examine the quality and contiguity of the scaffolded assembly, the assembly was aligned to a reference *A. thaliana* genome in protein space using promer from the mummer bioinformatic tool suite. The resulting plot displayed a mirroring pattern whereby scaffolds had apparent end-to-end joining of duplicate chromosomes (Supplemental Figure 1). This mirroring pattern suggested the possibility of two different phenomena occurring. The first was a potential whole genome duplication due to the frequency and size of the mirrored regions and the second was the presence of mis-joins in the assembly given the proximity of the mirrored regions to their counterparts. While the latter was deemed to be most likely a scaffolding artifact, the former was found to be biologically plausible given the frequency of recent whole genome duplications found across the different tribes of the Brassicaceae family (Kagale et al., 2014; Mandáková et al., 2017).

### Round 3: HiFi Assembly

To further investigate the possibility of a whole genome duplication and correct mis-joins created in the Hi-C scaffolding process, a third round of sequencing was performed. Leaf tissue was collected from individuals located in the same geographic region as the first two rounds and extracted high molecular weight DNA was prepped and sequenced on the PacBio Revio sequencing platform. Following postprocessing, the HiFi reads were assembled using the long read assembler HiFiasm. The HiFi assembly was an improvement over the assemblies created in both rounds one and two. Total sequence length increased to 402 Mb with 86.0% of the total sequence length contained in the 18 scaffolds whose length was greater than 1 Mb. Along with the increase in assembly size, the total number of scaffolds dropped to 1,193 and the scaffold N50 increased to 23.5 Mb (Table 1). The HiFi assembly also saw an improvement in complete BUSCOs and now showed slightly more duplicated than single-copy BUSCOs. All BUSCOs were also contained within the previously mentioned 18 scaffolds (Table 2). The eleven largest scaffolds had telomere repeats at both ends, indicating that these scaffolds likely represent telomere-to-telomere chromosome assemblies (Supplemental Figure 2). Of the remaining seven scaffolds larger than 1 Mb, six had telomere repeats at a single end and one had no repeats. Thus, the expected number of 28 telomeres was represented on the eighteen largest scaffolds. When compared to the round one assembly, most of the sequence is shared between the two assemblies, with the main difference being the contiguity of scaffolds (Figure 1).

**Figure 1.**
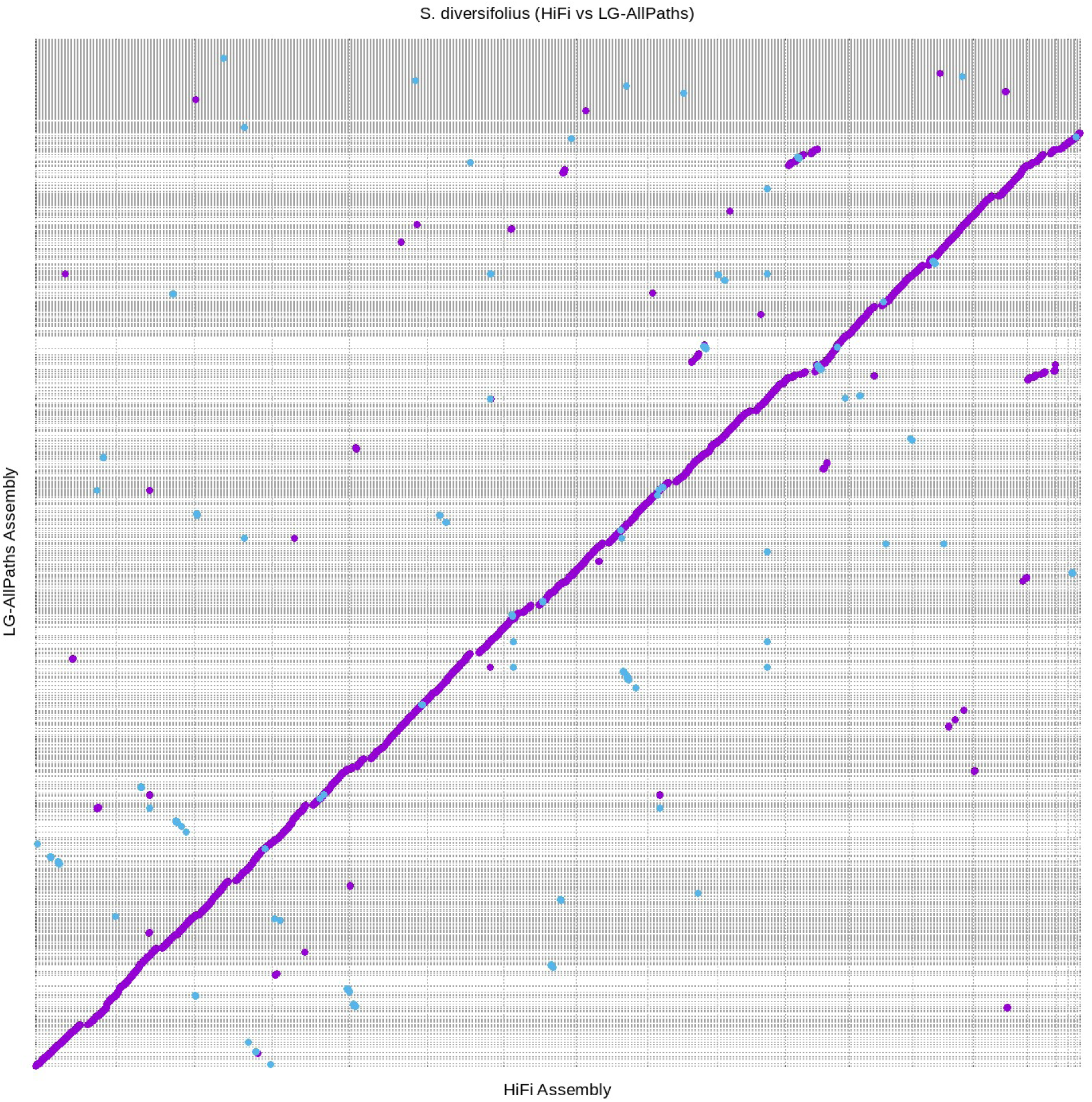
All scaffolds greater than 1 Mb in the HiFiasm assembly aligned to all scaffolds in the LG-AllPaths assembly. Alignment was performed using Nucmer and plotted using Mummerplot, both programs included in the Mummer bioinformatic toolkit. Purple lines indicate regions of alignment in the forward direction and blue lines indicate regions of alignment in the reverse direction. Gray dashed lines indicate separate scaffold in each assembly.

Annotation was completed using the Maker-P pipeline. Maker analysis of the HiFi assembly from round 3 predicted 40,606 protein-coding genes after filtering. Of these 40,606 protein-coding genes, 34,866 are found in the 18 largest scaffolds.

### Comparison of Round 2 and 3

The Hifi assembly was created to correct mis-joins created in the Hi-C scaffolding process and validate the existence of a recent whole genome duplication. When aligned against one another, the Hi-C scaffolded and HiFi assembly were mostly congruent. (Figure 2). The HiFi assembly was aligned to *A. thaliana* in the same way as the Hi-C scaffolded assembly. The mirroring pattern observed in the Hi-C scaffolded assembly was no longer present. Evidence of a recent whole genome duplication was present as seen by the presence of duplicate ancestral genomic regions spread throughout the genome (Figure 3).

**Figure 2.**
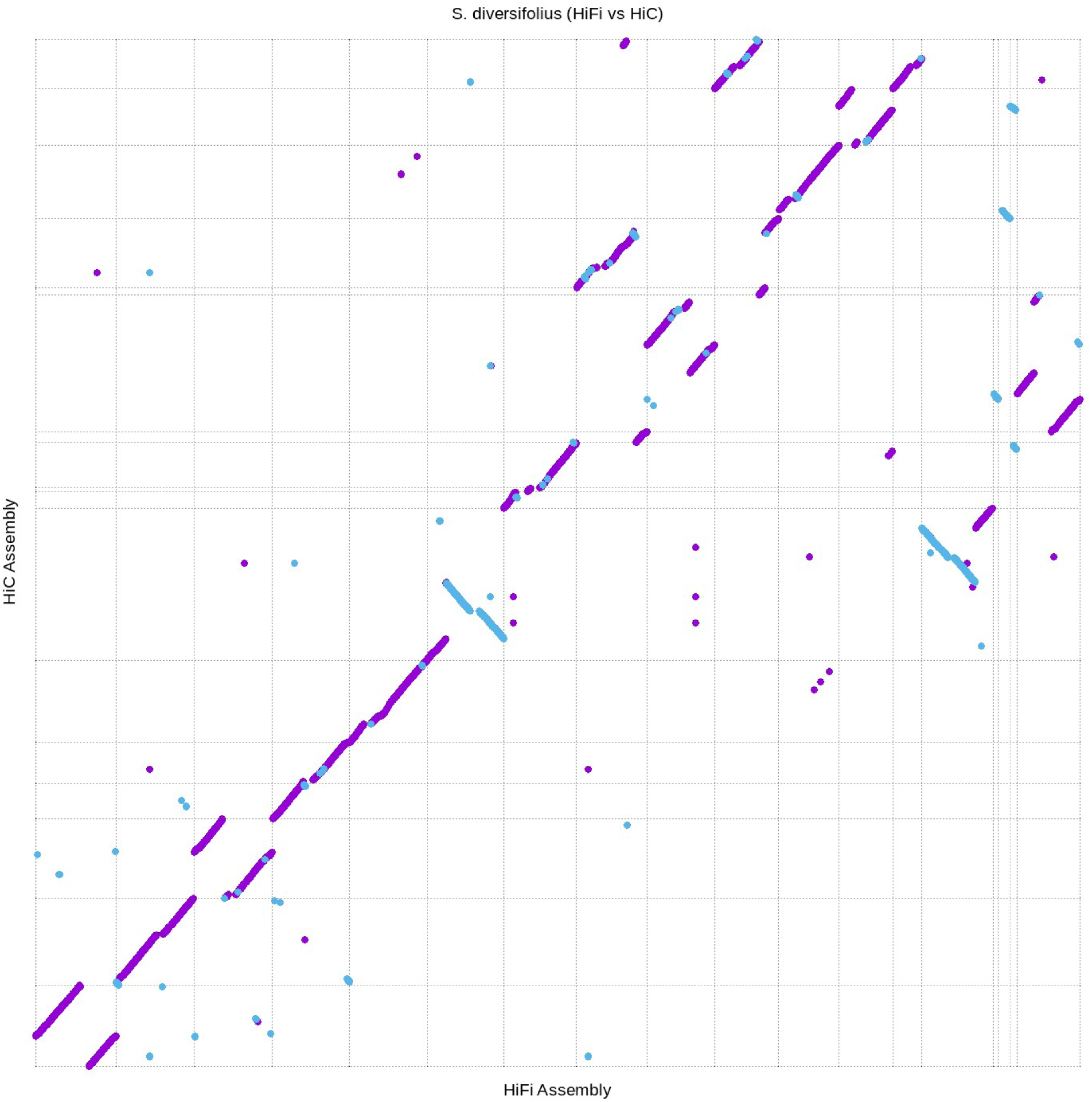
All scaffolds greater than 1 Mb in the HiFiasm assembly aligned to all scaffolds greater than 1 Mb in the HiC assembly. Alignment was performed using Nucmer and plotted using Mummerplot, both programs included in the Mummer bioinformatic toolkit. Purple lines indicate regions of alignment in the forward direction and blue lines indicate regions of alignment in the reverse direction. Gray dashed lines indicate separate scaffold in each assembly.

**Figure 3.**
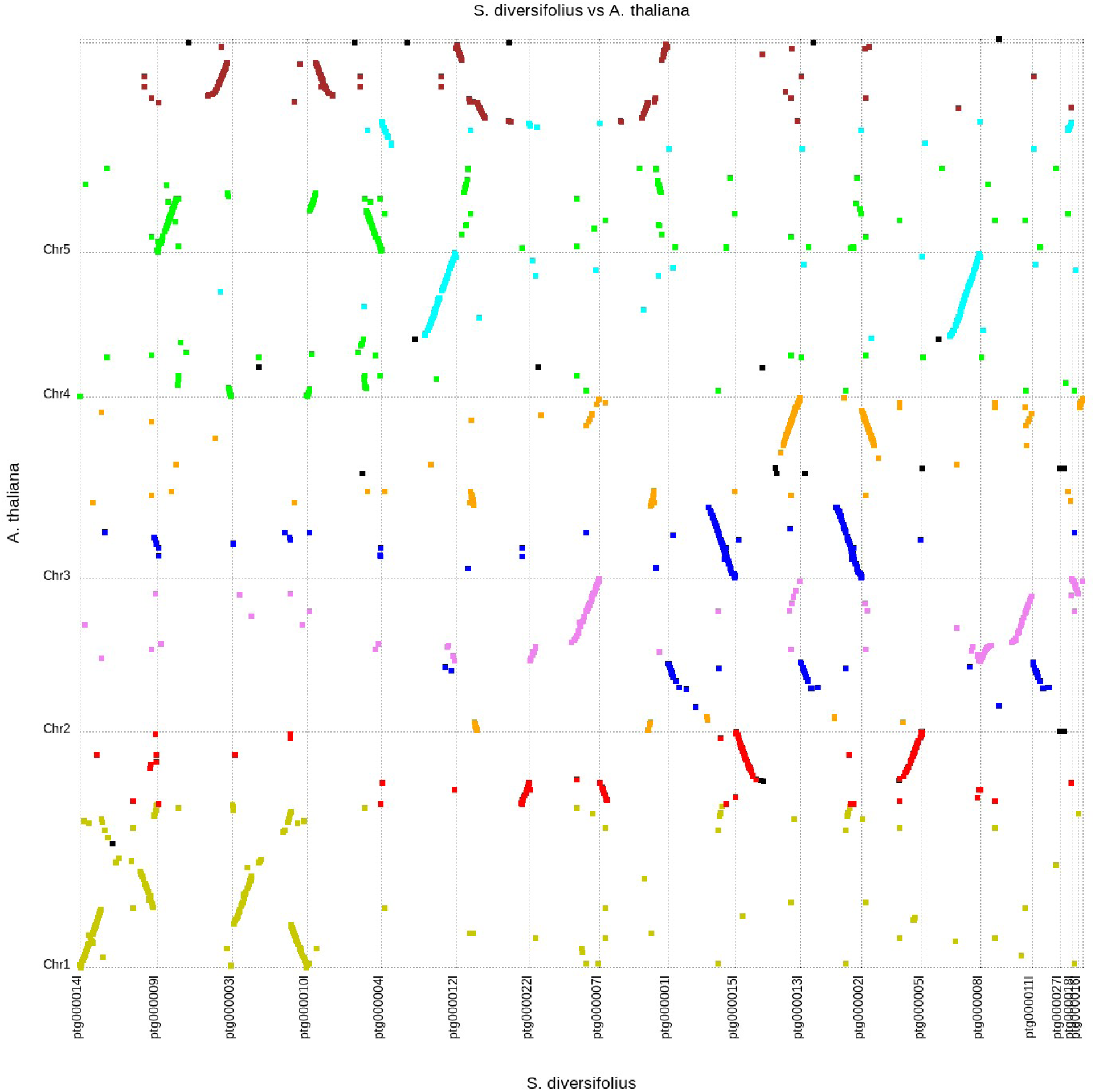
All scaffolds greater than 1 Mb in the HiFiasm assembly aligned to the TAIR 10 A. thaliana genome assembly. Alignment was performed using Promer and plotted using Mummerplot, both programs included in the Mummer bioinformatic toolkit. The color of each line corresponds to the 8 ancestral crucifer karyotype (ACK) blocks of A. thaliana with the exception of black which corresponds to regions which do not belong to an ACK block. ACK block boundaries are based on gene locations summarized in Lysak et. Al 2016. Gray dashed lines indicate separate scaffold in each assembly.

### Whole Genome Duplication

Our genome alignment plot provides additional evidence of a whole genome duplication shared among members of the Thelypodieae tribe. To investigate this further, we examined the distribution of the synonymous substitution rate (Ks) among paralogs. In the absence of whole genome duplication events, the Ks distribution is expected to show a rapid decrease over time (Lynch and Conery, 2000). Additional peaks of in the Ks distribution indicate whole genome duplication events and can be used to estimate their age (Maere et al., 2005; Ohno, 1970) Analysis of the Ks peaks for internal WGD gene pairs in *S. diversifolius* and the previously mentioned genomes revealed that *S. diversifolius* has a relatively new WGD with a Ks of ∼0.18, along with two older peaks at 0.92 and 2.03, likely corresponding to the Brassicaceae α WGD and the Eudicots γ WGT, respectively (Figure 4). Our constructed phylogenetic tree (Figure 5) which agrees with other phylogenetic analyses ((Ivalú Cacho et al., 2014; Kagale et al., 2014; Mandáková et al., 2017) suggests *S. diversifolius, C. amplexicaulis*, and *S. pinnata* are closely related and both *C. amplexicaulis* and *S. pinnata* also exhibit a Ks peak around 0.18. No other species in our analysis shared the Ks peak around 0.18. To determine the species divergence time and ascertain if it is the same WGD we calculated the Ks values between *S. diversifolius* and *S. pinnata* and between *C. amplexicaulis* and *S. diversifolius*. These species show a divergence Ks peak of about 0.08, which is later than the WGD event. Therefore, it appears this WGD, likely occurring around 15 MYA, is shared by *S. diversifolius* and other members of the Thelypodieae.

**Figure 4.**
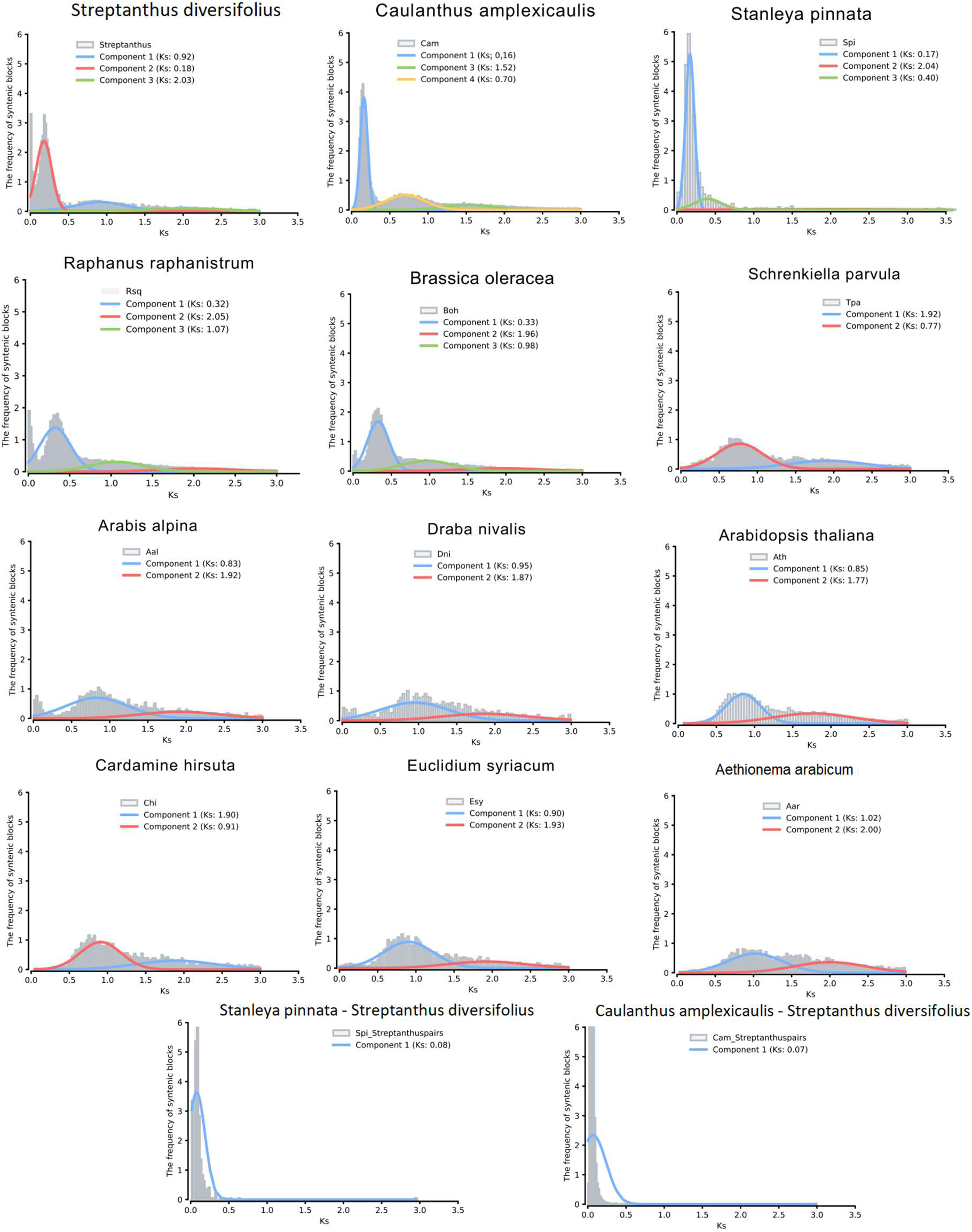
Top plots are intragenomic analysis of Ks for the analyzed species (excluding T. arvense and A. Lyrate). Colored lines indicated distinct Ks peaks in each plot. Bottom plots are intergenomic analysis of Ks between S. diversifolius and either S. pinnata or C. amplexicaulis

**Figure 5.**
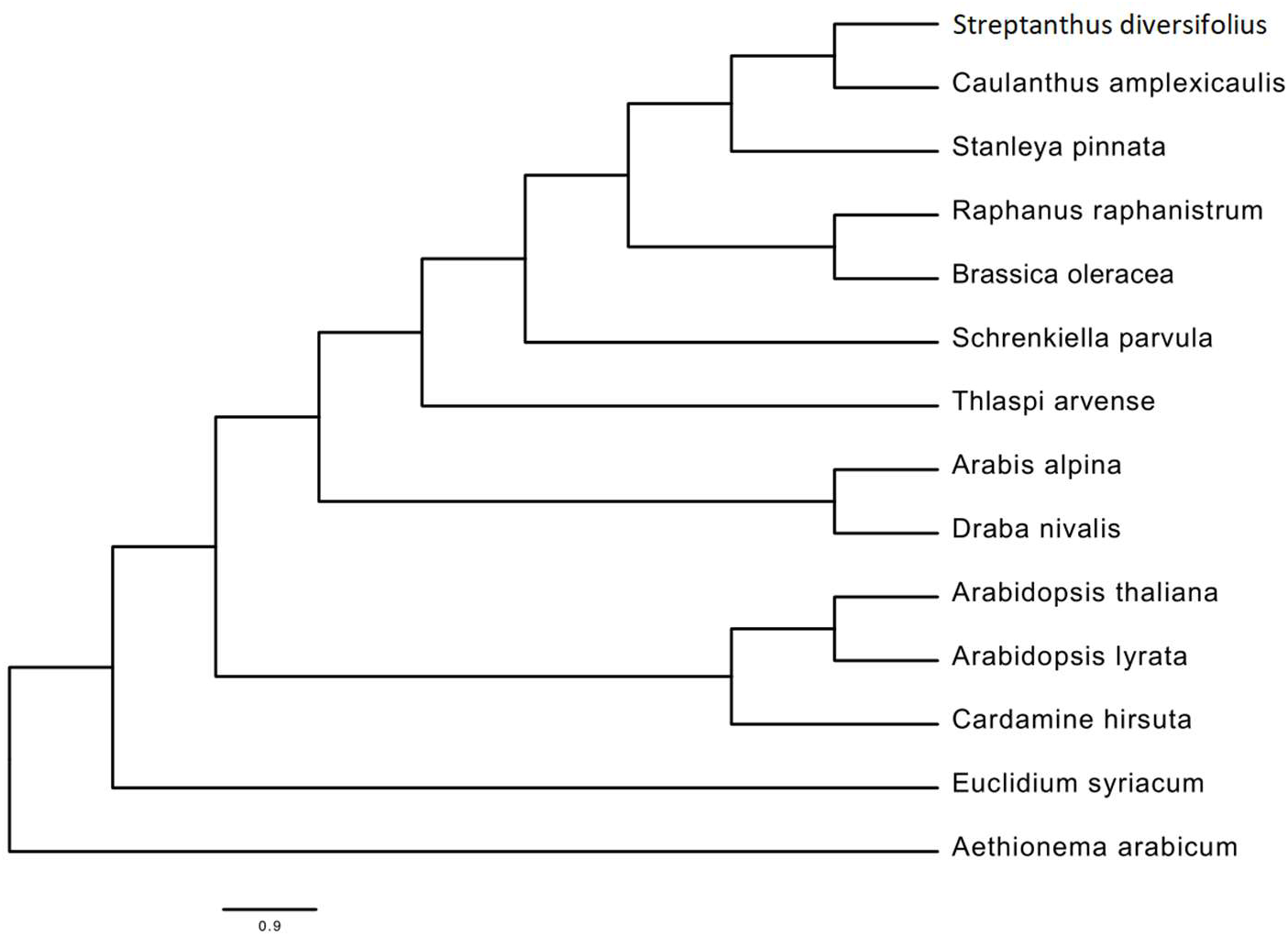
Phylogenetic species tree. Tree was inferred using phylogenetic trees from the 118 single-copy orthogroups identified between our 14 species of interest.

Whole genome duplication can result from non-disjunction during meiosis (creating an autotetraploid) or by hybridization of closely related species (creating an allotetraploid). We aimed to distinguish these two possibilities by comparing gene and repeat content of homoeologous chromosomes. It is well documented that genes are retained unequally after allotetraploidization, with one sub-genome becoming more fractionated than the other (Freeling et al., 2012; Sankoff et al., 2010; Thomas et al., 2006). We performed a fractionation bias analysis using CoGe (Joyce et al., 2017; Lyons and Freeling, 2008) to compare gene loss of homoeologous *Strepanthus diversifolius* scaffolds, relative to *Arabidopsis thaliana.* We found no evidence of fractionation bias (Figure 6). For relatively recent allotetraploids, one also expects two find homoeolog-specific repeat sequences, because the repeat sequences will have diverged over evolutionary time in the two progenitors. To search for sub-genome specific repeats, we used SubPhaser (Jia et al., 2022) to look for kmers that could distinguish between homeologs. Of 4,425,565 kmers identified, only 88 were unique to one homeolog among a pair, and there was no significant enrichment of unique kmers on any homeolog. Taken together, these analyses suggest that the whole genome duplication described here either resulted from an autopolyploid event. or from hybridization of two very closely related species.

**Figure 6.**
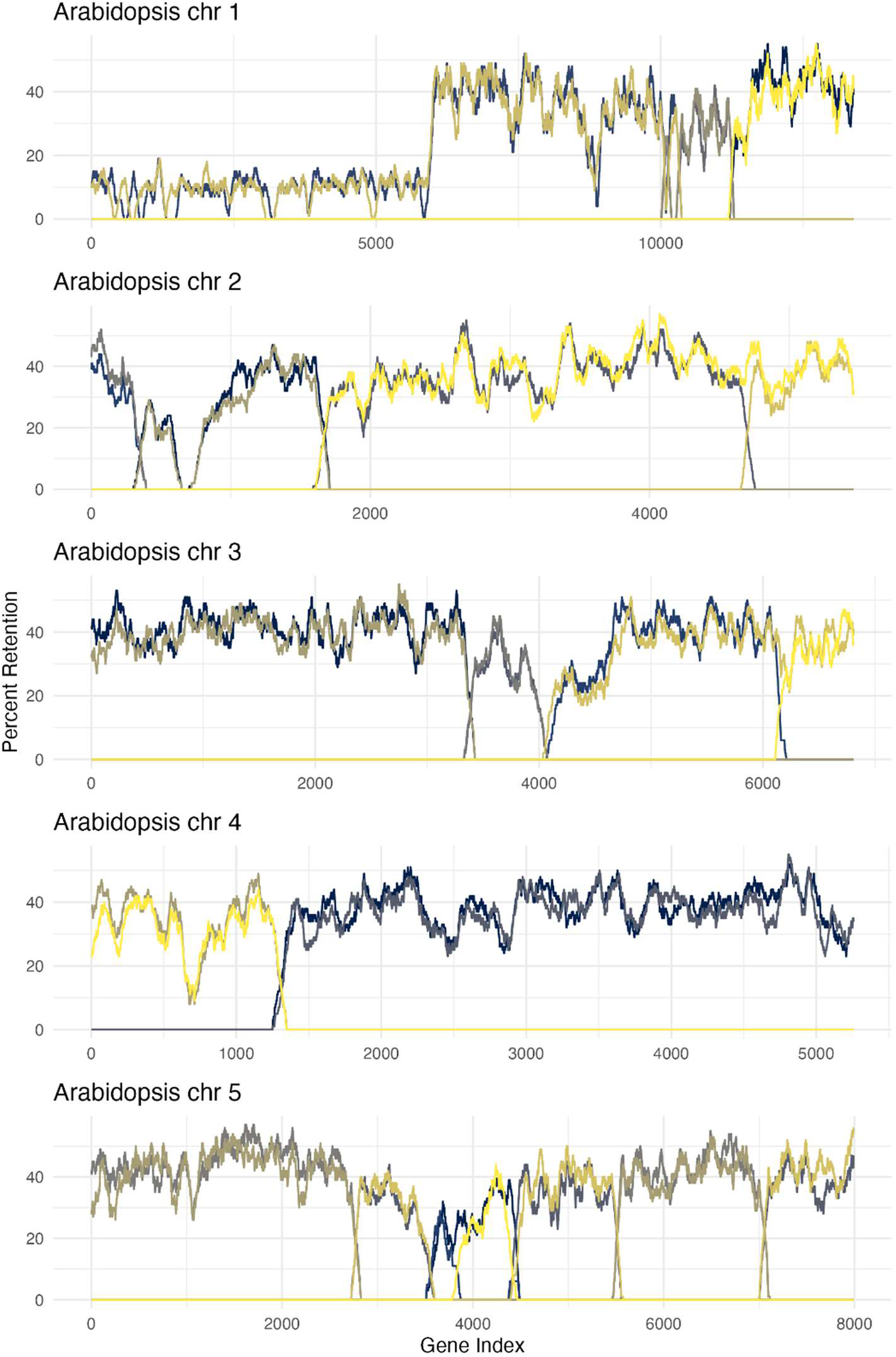
*Fractionation Bias*. The plot shows the percent gene retention of each *S. diversifolius* scaffold (colored lines) when compared to *Arabidopsis thaliana*. Colors are not conserved across different *Arabidopsis* chromosomes (i.e. yellow on chr 1 plot may represent a different *S. diversifolius* scaffold than yellow on chr 2).

### Gene family expansion and duplicate gene retention

Confident in the presence of a WGD we further looked at the composition of the genome to assess if there were gene families that showed significant expansion, or gene categories with preferential retention of both duplicates. CAFE5 analysis reported a total of 19 expanded gene families containing a total of 295 genes (Figure 7). GO and KEGG enrichment analysis were performed on this set of genes. GO analysis revealed an enrichment for several biological processes related to ATP. KEGG analysis also showed an enrichment for ATP related processes with the largest most significant category being F-type H+/Na+-transporting ATPase subunit alpha-EC:7.1.2.2 7.2.2.1. The enrichment of ATP related terms is of interest given serpentine soil habitats occupied by many *Streptanthus* species are characterized as having higher levels of magnesium along with elevated levels of other heavy metal which are toxic to most plants (Brady et al., 2005). Chelation of nucleotides by magnesium (Mg) is an essential feature of cell metabolism with adenylates being the most abundant (Kleczkowski and Igamberdiev, 2021). The excess Mg of the serpentine soil needs to be managed to maintain a [Mg^2+^] that allows the plant to perform metabolic tasks without damage from Mg^2+^ toxicity. One way that plants avoid Mg^2+^ toxicity is by utilizing the CBL-CIPK network which allows cells to sequester excess Mg^2+^ into vacuoles (Tang et al., 2015).

**Figure 7.**
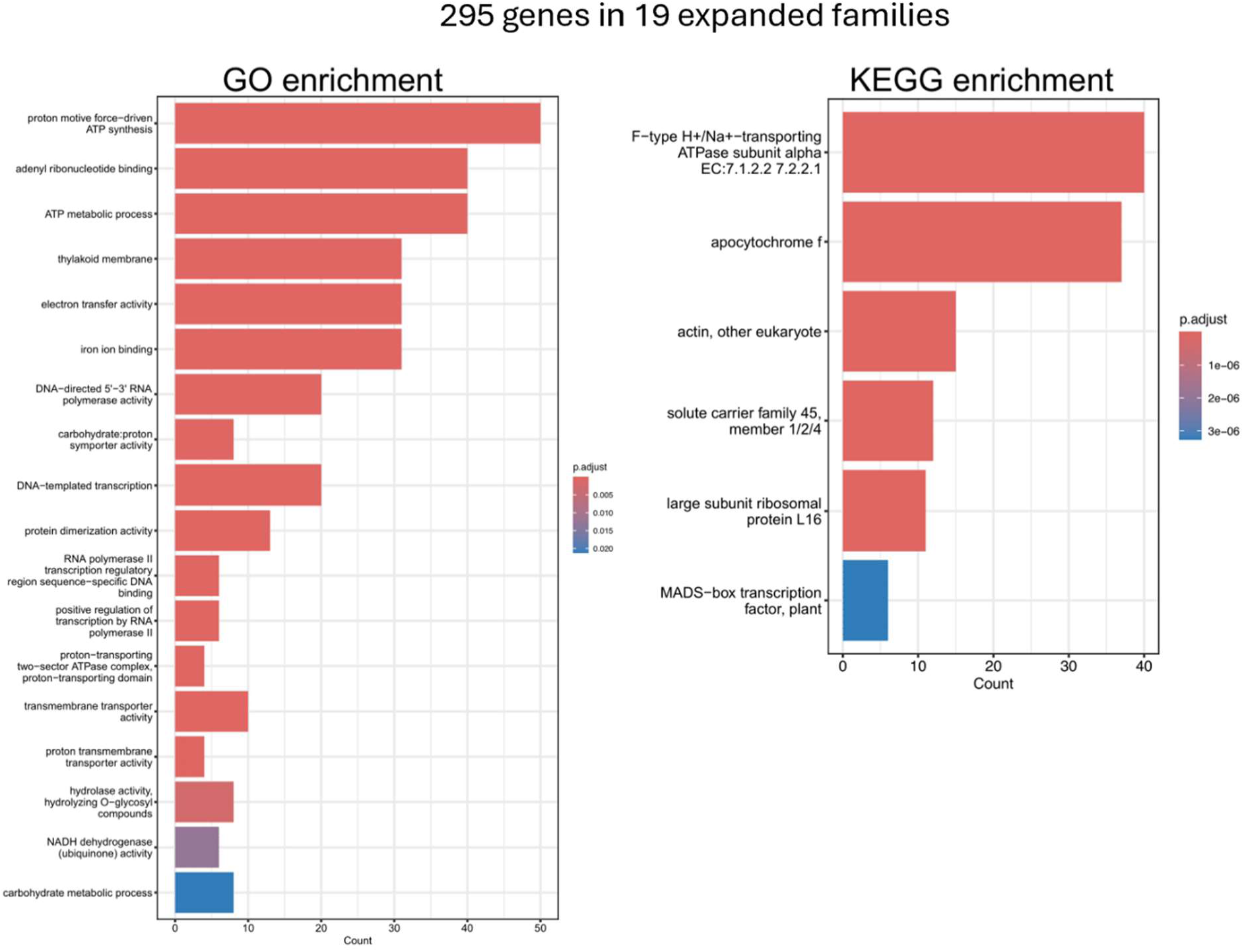
GO and KEGG enrichment analysis of the 295 genes in the 19 expanded gene families.

Along with enrichment among gene families, GO and KEGG enrichment analysis were also performed on WGD-derived gene pairs identified through the DupGen_finder pipeline. Of note, KEGG analysis identified 3 categories significantly enriched including calcium-dependent protein kinase EC:2.7.11.1 (Figure 8). Enrichment of these genes may be benefiting the plant as it negotiates a Mg rich environment potentially by being involved in the CBL-CIPK network or improving signaling to maintain appropriate [Mg^2+^] in the cell.

**Figure 8.**
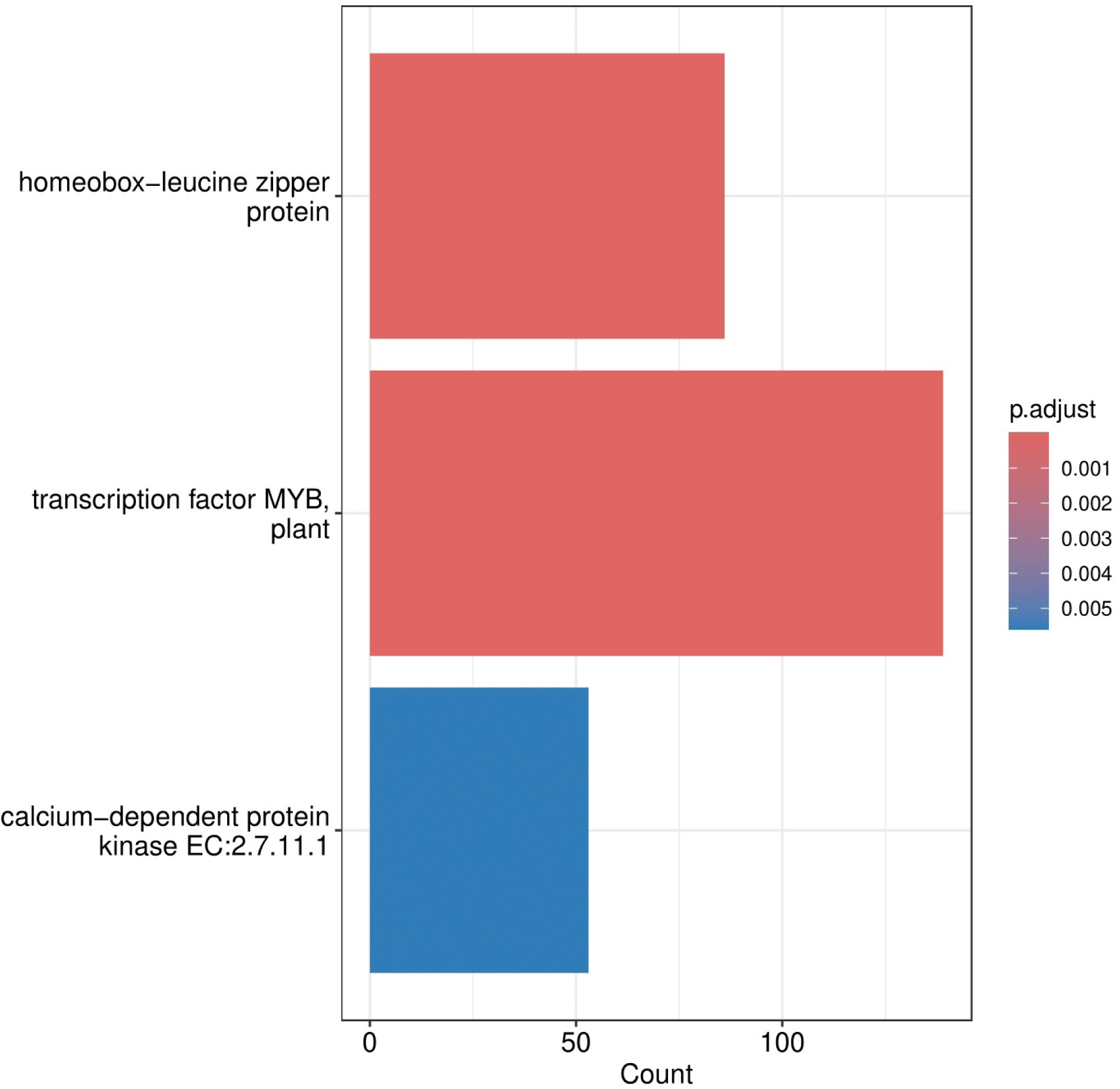
KEGG enrichment analysis of WGD-derived gene pairs. WGD-derived genes identified through GenDup_finder pipeline

### Positive selection of duplicated genes

Analysis of the whole genome duplicated genes found a total of 123 Streptanthus genes to be under positive selection (Supplemental Table 2). However, there was no significant KEGG or GO enrichment after false-discovery rate correction.

## Discussion

The Streptanthoid Complex is a collection of plants found throughout the California Floristic Province. Individuals in this collection can be found in environments ranging from arid deserts all the way to snow packed mountain tops (Baldwin et al., 2012). Despite living in drastically different environments, these flowering plants share a highly connected genetic background. Phylogenetically close relationships along with the expansive range and diverse morphology make this collection of plants an ideal study subject for exploring the rapid radiation and adaptation of flowering plants in the California Floristic Province. Active research is currently being conducted on this complex (Cacho et al., 2021, 2015; Cacho and Strauss, 2014; Gremer et al., 2020; Weber et al., 2018) but is restricted to variation in traits due to a lack of genomic resources. This assembly offers a starting point in expanding the realm of genomic analyses related to these species. Future studies will now have another tool to help bridge the gap between the observed phenotypes and the underlying genetics.

The assembly presented here provides further evidence of a recent whole genome duplication shared throughout the Thelypodieae tribe which contains the Streptanthoid Complex. Whole genome duplications have previously been described in other members of the tribe, including *S. farnthworthianus* (Mandáková et al., 2017), *C. amplexicaulis* (Burrell et al., 2011), *P. antiscorbutica* and *Stanleya pinnata* (Kagale et al., 2014). Prior studies used transcriptome or molecular marker analysis. We build on these studies by providing a whole genome assembly that can be used to study genome evolution after the whole genome duplication, and through our cross-species Ks analysis that indicates there was a single, shared WGD event among these species.

Gene duplication is a critical component of evolution, allowing rapid evolution of new functions (Ohno, 1970). Following a whole genome duplication event, there is typically a rapid extensive process of genome rearrangement and consolidation as diploidization occurs (Lynch and Conery, 2000; Qiao et al., 2019). During this time, genes and their functions can be lost, gained, and changed through processes including subfunctionalization and neofunctionalization (Almeida-Silva and Van de Peer, 2023; Lynch and Conery, 2000; Qiao et al., 2019). It has been reported that the At-β whole genome duplication event shared among the order Brassicales expanded the ability of plants in this order to produce glucosinolates (Barco and Clay, 2019). It is possible that the recent whole genome duplication described here played a key role in rapid radiation of the Streptanthoid Complex throughout California. *Streptanthus* species may have been able to rapidly adapt to diverse and harsh environments such as serpentine soils through the increased genetic arsenal following the whole genome duplication (Ohno, 1970; Qiao et al., 2019). For example, it is possible that the expansion of genes associated with ATP related processes and calcium signaling has developed a secondary regulatory network able to handle excess external Mg. This would have allowed *Streptanthus* to occupy previously unexploited habitats including stressful serpentine soils characterized by Mg toxicity, and low Ca:Mg ratios. Future research should be directed towards identifying genes that allowed *Streptanthus* to expand its geographic and climatic range. Identification of these genes could have major impacts on the agricultural field given the close genetic relationship between *Streptanthus* and it economically important sister clade *Brassica*.

## Data Availability Statement

Genome sequencing data and assembly data can be found on NCBI under Bioproject PRJNA283414

## Acknowledgments

We thank the Department of Energy Joint Genome Institute and collaborators for prepublication access to the *Caulanthus amplexicaulis* and *Stanleya pinnata* genome sequences. The work (proposal: 10.46936/10.25585/60000980) conducted by the U.S. Department of Energy Joint Genome Institute (https://ror.org/04xm1d337), a DOE Office of Science User Facility, is supported by the Office of Science of the U.S. Department of Energy operated under Contract No. DE-AC02-05CH11231. This work used Jetstream2 (Hancock et al., 2021) at Indiana University through allocation BIO230019 from the **<u>Advanced Cyberinfrastructure</u> <u>Coordination Ecosystem: Services & Support</u>** (ACCESS) program (Boerner et al., 2023), which is supported by National Science Foundation grants #2138259, #2138286, #2138307, #2137603, and #2138296. We thank Dylan Burge for participating in project initiation, sample collection, DNA isolations, and sequencing for the initial Illumina assembly.

## Conflict of Interest

The authors report no conflict of interest.

## Funder Information

Natural Sciences and Engineering Research Council of Canada, Grant ID 327426, to LHR. United States National Science Foundation grant 1831913 to JRG, JNM, SYS, JTD. The National Institute of Food and Agriculture, United States Department of Agriculture (USDA-NIFA) award CA-D-PLB-2795-H to JNM. China Scholarship Council Scholarship grant 202006850062 to QL.

## Supplemental figures

**Supplemental Figure 1.**
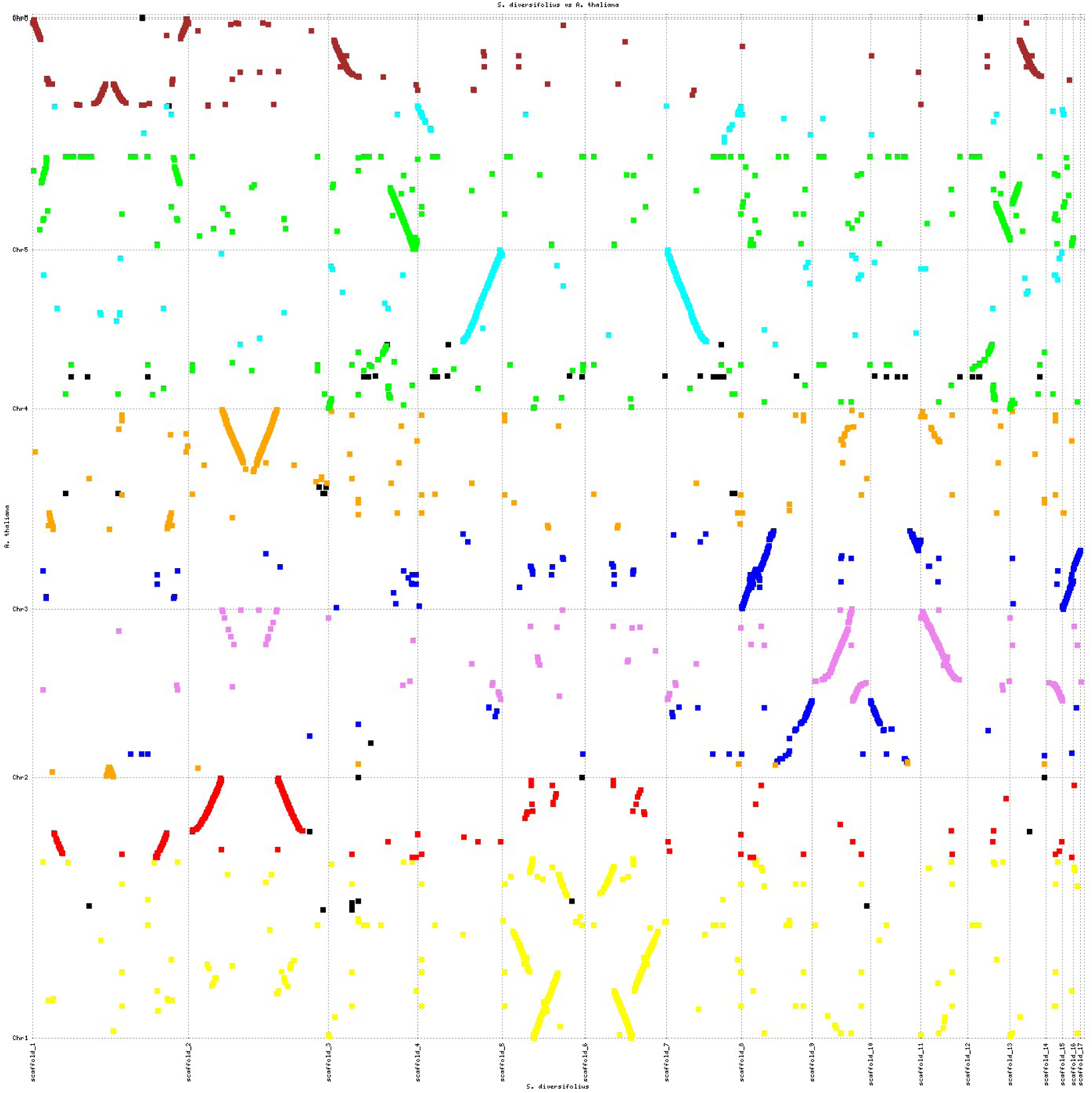
All scaffolds greater than 1 Mb in the HiC assembly aligned to the TAIR 10 A. thaliana genome assembly. Alignment was performed using Promer and plotted using Mummerplot, both programs included in the Mummer bioinformatic toolkit. The color of each line corresponds to the 8 ancestral crucifer karyotype (ACK) blocks of A. thaliana with the exception of black which corresponds to regions which do not belong to an ACK block. ACK block boundaries are based on gene locations summarized in Lysak et. Al 2016. Gray dashed lines indicate separate scaffold in each assembly.

**Supplemental Figure 2.**
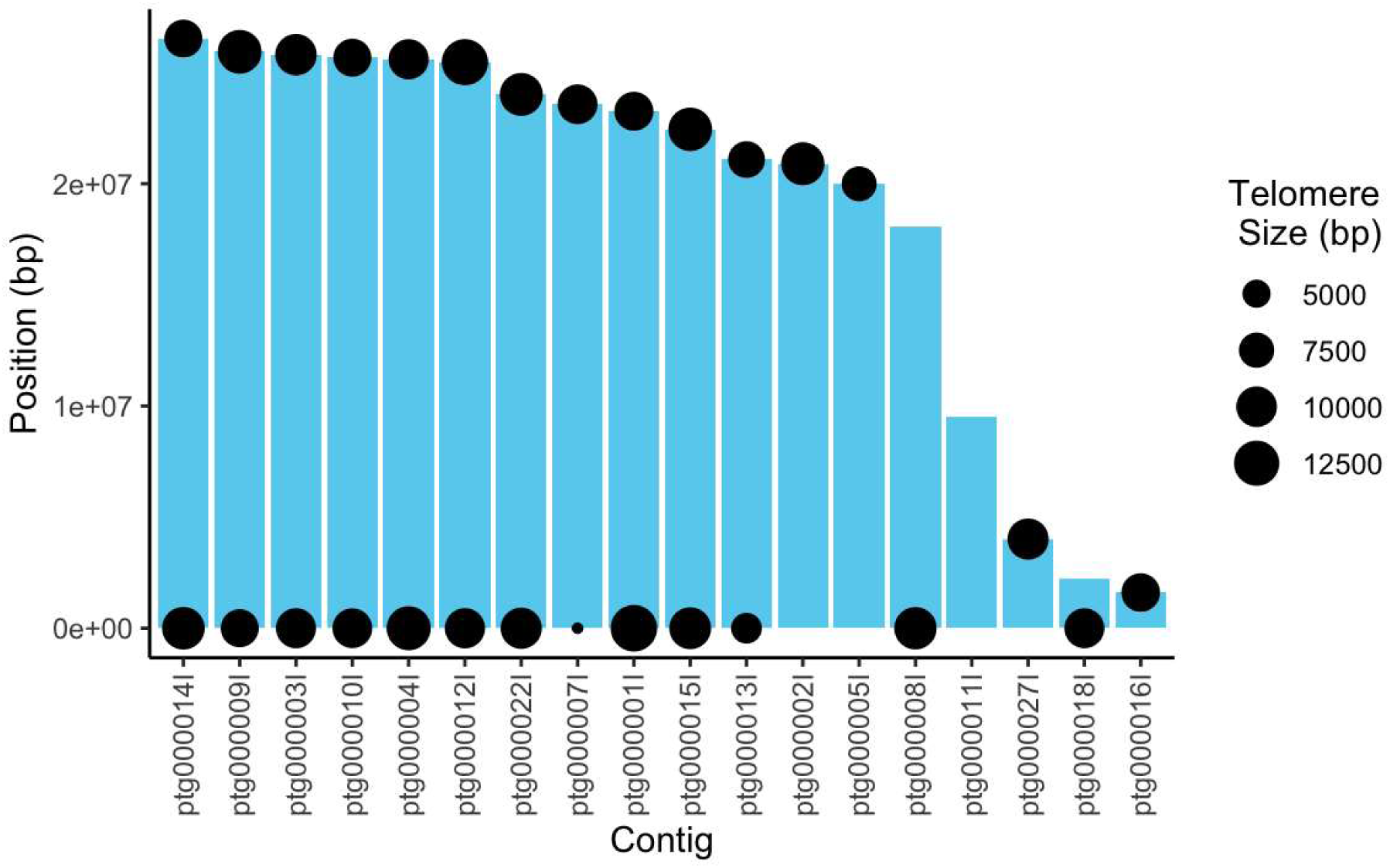
Illustration of telomere repeat location on the 18 scaffolds larger than 1 Mb.

## Supplemental tables

**Supplemental Table 1.** Repeat element content of HiFi assembly. Analysis completed using RepeatMasker version 4.1.5 in sensitive mode. Query species was assumed to be Brassicales.

|  |  |  | Number<br>of<br>elements | Length<br>occupied (bp) | percentage of<br>sequence (%) |
| --- | --- | --- | --- | --- | --- |
| **Retroelements** |  |  | 74,066 | 75,569,968 | 18.79 |
|  | SINEs: |  | 270 | 39,770 | 0.01 |
|  | Penelope: |  | 0 | 0 | 0 |
|  | LINEs: |  | 4,782 | 3,064,392 | 0.76 |
|  |  | CRE/SLACS | 0 | 0 | 0 |
|  |  | L2/CR1/Rex | 0 | 0 | 0 |
|  |  | R1/LOA/Jockey | 0 | 0 | 0 |
|  |  | R2/R4/NeSL | 0 | 0 | 0 |
|  |  | RTE/Bov-B | 0 | 0 | 0 |
|  |  | L1/CIN4 | 4,758 | 3,061,849 | 0.76 |
|  | LTR elements |  | 69,014 | 72,465,806 | 18.02 |
|  |  | BEL/Pao | 0 | 0 | 0 |
|  |  | Ty1/Copia | 32,174 | 42,611,770 | 10.6 |
|  |  | Gypsy/DIRS1 | 33,841 | 28,849,738 | 7.17 |
| **DNA transposons** |  |  | 20,796 | 6,705,897 | 1.67 |
|  | hobo-Activator |  | 4,574 | 1,292,104 | 0.32 |
|  | Tc1-IS630-Pogo |  | 1,476 | 311,487 | 0.08 |
|  | En-Spm |  | 0 | 0 | 0 |
|  | MULE-MuDR |  | 6,750 | 2,499,600 | 0.62 |
|  | PiggyBac |  | 0 | 0 | 0 |
|  | Tourist/Harbinger |  | 1,260 | 438,400 | 0.11 |
|  | Other (Mirage, P-element, Transib) |  | 0 | 0 | 0 |
| **Rolling-circles** |  |  | 994 | 159,073 | 0.04 |
| **Unclassified:** |  |  | 23 | 4,153 | 0 |
| **Total interspersed repeats:** |  |  |  | 82,280,018 | 20.46 |
| **Small RNA** |  |  | 7,919 | 12,239,420 | 3.04 |
| **Satellites:** |  |  | 58 | 10,415 | 0 |
| **Simple repeats:** |  |  | 120,687 | 16,567,216 | 4.12 |
| **Low complexity:** |  |  | 26,984 | 1,340,333 | 0.33 |

**Supplemental Table 2.**
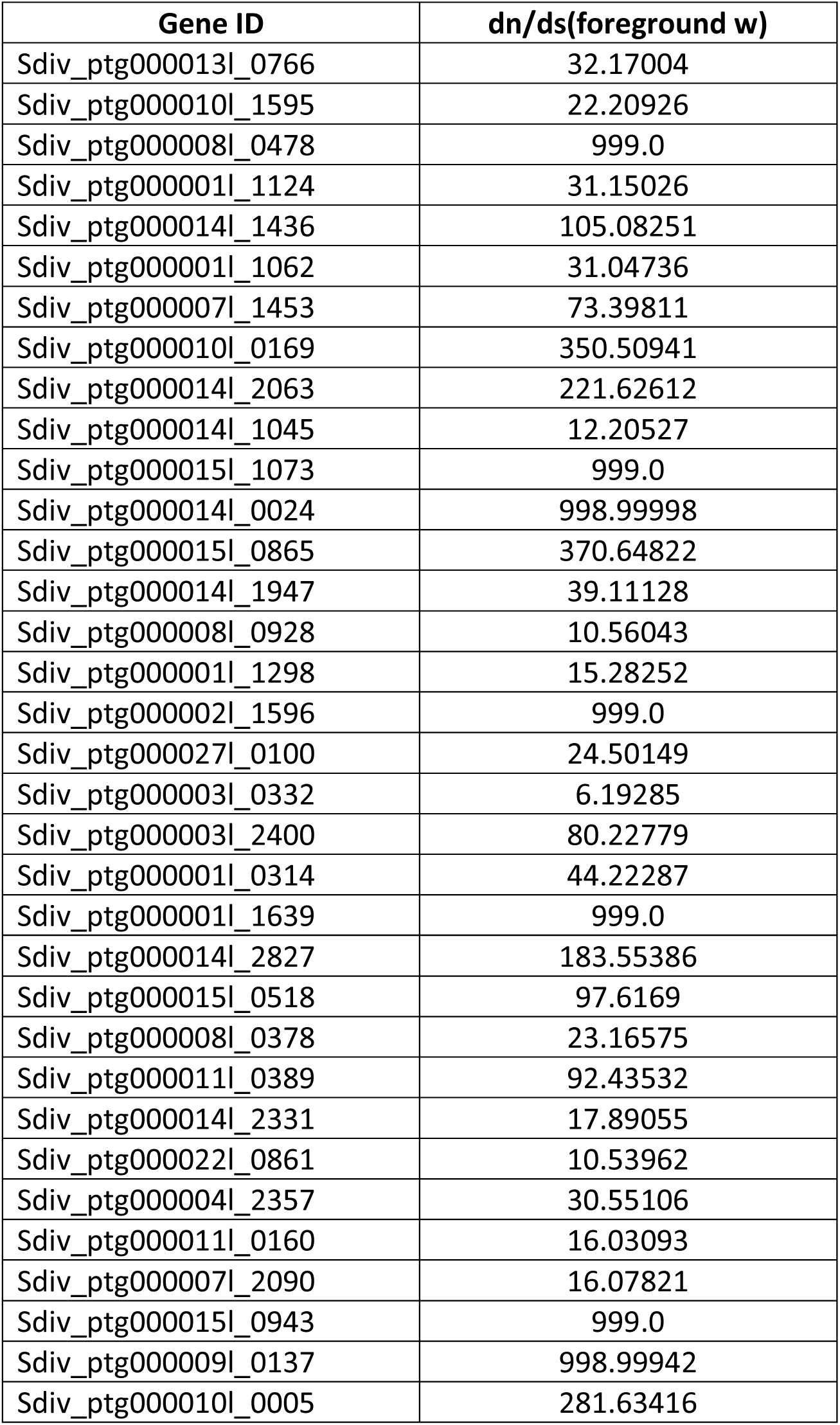
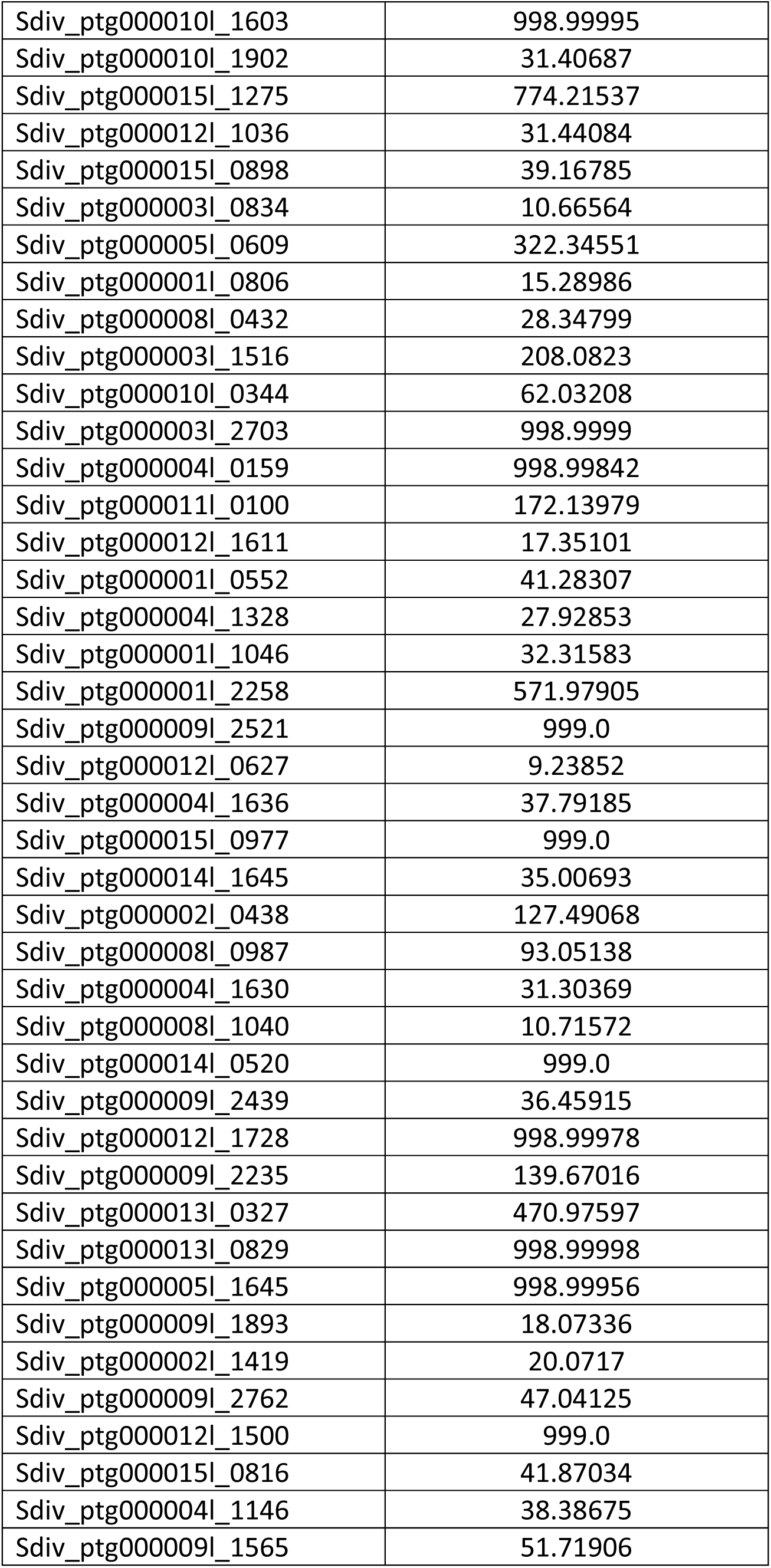
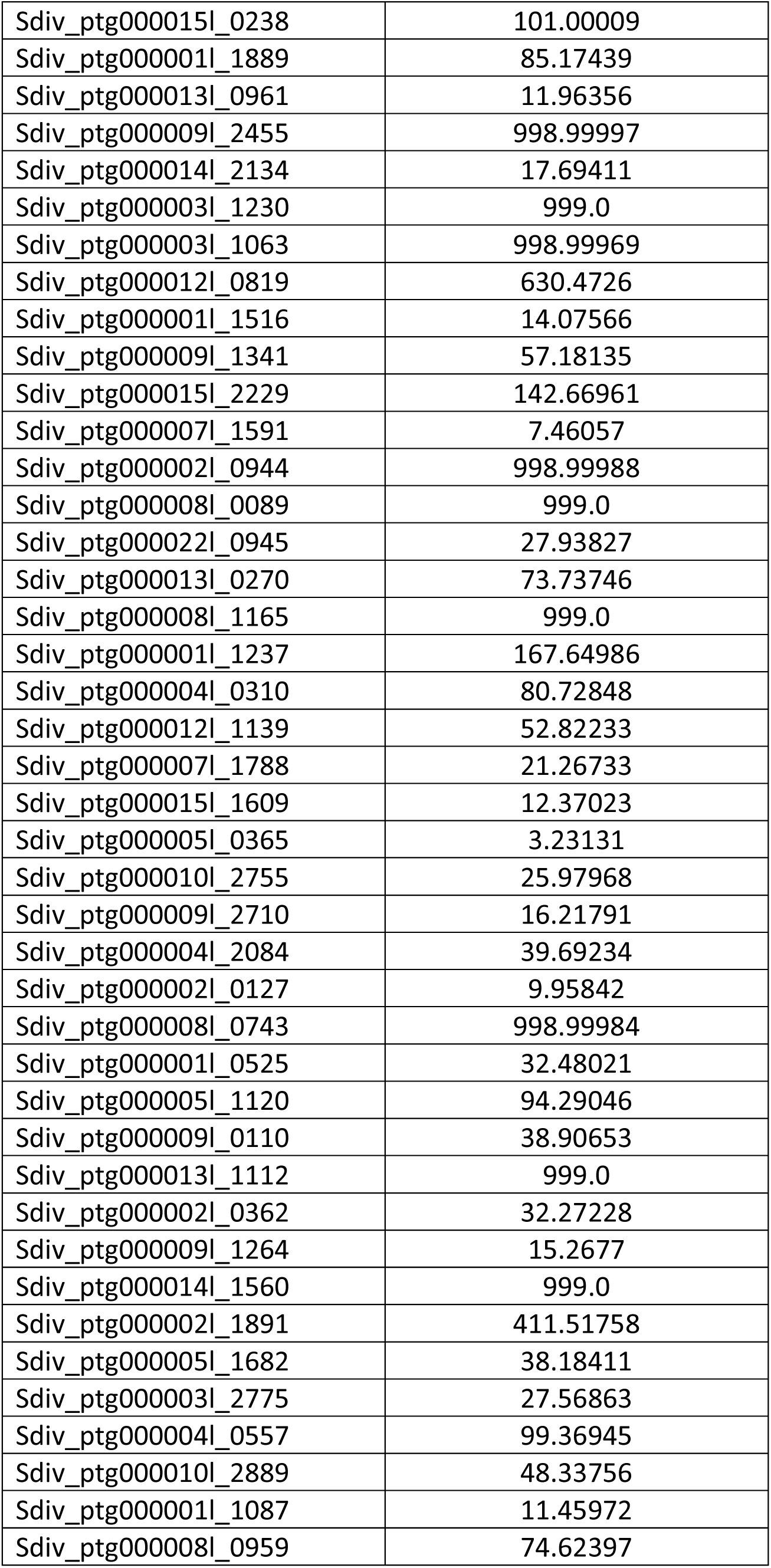
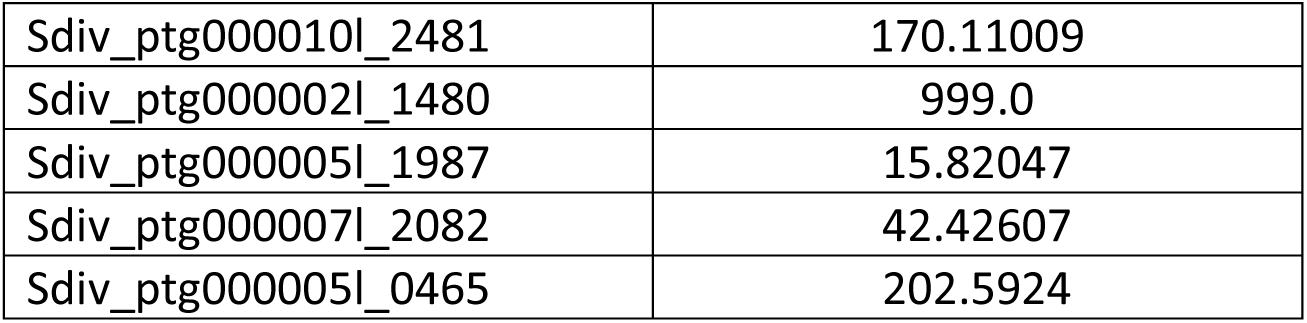
Whole genome duplicated genes identified as being positively selected.

